# Blocking Src-PSD-95 interaction rescues glutamatergic signaling dysregulation in schizophrenia

**DOI:** 10.1101/2024.03.08.584132

**Authors:** Robert E. Featherstone, Hongbin Li, Ameet Sengar, Karin E. Borgmann-Winter, Olya Melnychenko, Lindsey M. Crown, Ray L. Gifford, Felix Amirfathi, Anamika Banerjee, Krishna Parekh, Margaret Heller, Wenyu Zhang, Adam D. Marc, Michael W. Salter, Steven J. Siegel, Chang-Gyu Hahn

## Abstract

The complex and heterogeneous genetic architecture of schizophrenia inspires us to look beyond individual risk genes for therapeutic strategies and target their interactive dynamics and convergence. Postsynaptic NMDA receptor (NMDAR) complexes are a site of such convergence. Src kinase is a molecular hub of NMDAR function, and its protein interaction subnetwork is enriched for risk-genes and altered protein associations in schizophrenia. Previously, Src activity was found to be decreased in post-mortem studies of schizophrenia, contributing to NMDAR hypofunction. PSD-95 suppresses Src via interacting with its SH2 domain. Here, we devised a strategy to suppress the inhibition of Src by PSD-95 via employing a cell penetrating and Src activating PSD-95 inhibitory peptide (TAT-SAPIP). TAT-SAPIP selectively increased post-synaptic Src activity in humans and mice, and enhanced synaptic NMDAR currents in mice. Chronic ICV injection of TAT-SAPIP rescued deficits in trace fear conditioning in Src hypomorphic mice. We propose blockade of the Src-PSD-95 interaction as a proof of concept for the use of interfering peptides as a therapeutic strategy to reverse NMDAR hypofunction in schizophrenia and other illnesses.

---

All antipsychotics currently prescribed for schizophrenia employ the same strategy, blockade of dopaminergic receptor subtype 2 and/or 5HT2A receptors(*1*). These agents have been the mainstay therapeutics of schizophrenia for seven decades, while their limitations have been abundantly recognized(*2*). The recently emerging genetic architecture of schizophrenia, however, does not lend strong support for targeting these receptors or any individual genes(*3*). It is now clear that schizophrenia is caused by multitudes of risk genes, many of which are shared by other neuropsychiatric illnesses(*4, 5*), and yet these risk genes precipitate the illness via specific modes of interactions and convergence (*6–8*) Thus, investigations of pathophysiologic mechanisms should incorporate the interactive dynamics and convergence mechanisms as well as dysfunction of individual genes(*5, 9*). Presently, mechanistic substrates for interactions and convergence are unknown and have not been exploited as an avenue for new therapeutic strategies for schizophrenia.

Gene-gene interactions and their pathophysiologic implications are notoriously difficult to study in a whole biological system. It is possible, however, when the interactions are enclosed in a subcellular locale and driven by known and quantifiable mechanistic substrates. The postsynaptic density (PSD) at the glutamatergic synapse could be such a locale, where schizophrenia risk genes are most enriched(*10*) and impact the function of the synapse via their physical interactions(*11*). The susceptibility for schizophrenia conferred by schizophrenia risk genes is strikingly enhanced when assessed within their networks constructed based on protein-protein interactions(*6, 7, 12, 13*). This is particularly pronounced in the glutamatergic synapse, suggesting protein interactions as potential mechanistic substrates for the interactive dynamics between synaptic risk genes.

Our recent studies demonstrated that altered protein interactions in NMDAR complexes converge on and reduce the activity of the non-receptor tyrosine kinase, Src, and precipitate NMDAR hypofunction in schizophrenia (*13*). We found that the DLPFC of schizophrenia subjects exhibit a striking decrease in tyrosine phosphorylation of the GluN2 subunits, which is indicative of decreased NMDAR function in patients with schizophrenia. This functional dysregulation of NMDARs was found to be mediated by hypoactivity of Src. Src hypoactivity was not associated with decreased expression of Src or its interactors but was accompanied by altered protein interactions including increased association of Src with PSD-95 and erbB4 and decreased Src association with rPTPa and dysbindin1 (*13*). Each of these aberrant protein interactions can reduce Src activity(*13*) and subsequently impact NMDAR currents through phosphorylation of NR2 subunits.

We then asked If these altered protein associations could be leveraged to rescue NMDAR hypofunction. Of the protein interactions described above, the Src-PSD-95 interaction is of particular interest as a potential therapeutic target since PSD-95 is increased in NMDAR complexes in schizophrenia and PSD-95 inhibits Src activity through direct interaction with the SH2 domain of Src(*14*). Here, we designed and tested a strategy to selectively target and reduce the interaction between Src and PSD-95, and thereby to selectively enhance Src activity in the NMDAR complexes. To enhance Src activity, we employed a Src activating PSD-95 inhibitory peptide (SAPIP) comprising the Src SH2 domain with a R175K mutation to minimize phosphotyrosine binding(*15*). SAPIP prevents PSD-95 from binding to Src, thereby overcoming inhibition of Src activity(*15*). This study tests the hypotheses that an interfering peptide intervention in the PSD-95–Src association will; 1) rescue NMDAR hypofunction *in vitro as assessed with NMDAR phosphorylation and protein-protein binding*, 2) normalize *ex vivo* synaptic currents and 3) improve cognitive behavioral and electrophysiological phenotypes *in vivo* in mice.

## Results

### TAT-SAPIP crosses cell membranes and enhances Src activity by blocking binging to PSD-95

To develop a form of SAPIP that may enter cells *in situ*, while retaining its activity, we fused the membrane-transduction sequence of HIV-TAT to the N-terminus of SAPIP (TAT-SAPIP). We first examined if TAT-SAPIP penetrates cell membranes and decreases Src–PSD-95 association. Primary rat cortical neurons were treated with TAT-SAPIP in the presence or absence of NMDA (10 uM) and glycine (1 uM). Synaptosomes from these cells were analyzed for protein associations between Src and PSD-95 using immunoprecipitation. We found significantly decreased PSD-95-Src association at 60 nM and 300 nM (F(2,12)=7.46, p= 0.012, 0.019 respectively) in the absence of NMDA receptor activation, as well as in the presence of it (F(2,12)=7.919, p=0.072, 0.005, respectively) (Figure 1 A,B). This suggests that TAT-SAPIP not only penetrates the cell membrane but also decreases Src–PSD-95 association.

**Figure 1:**
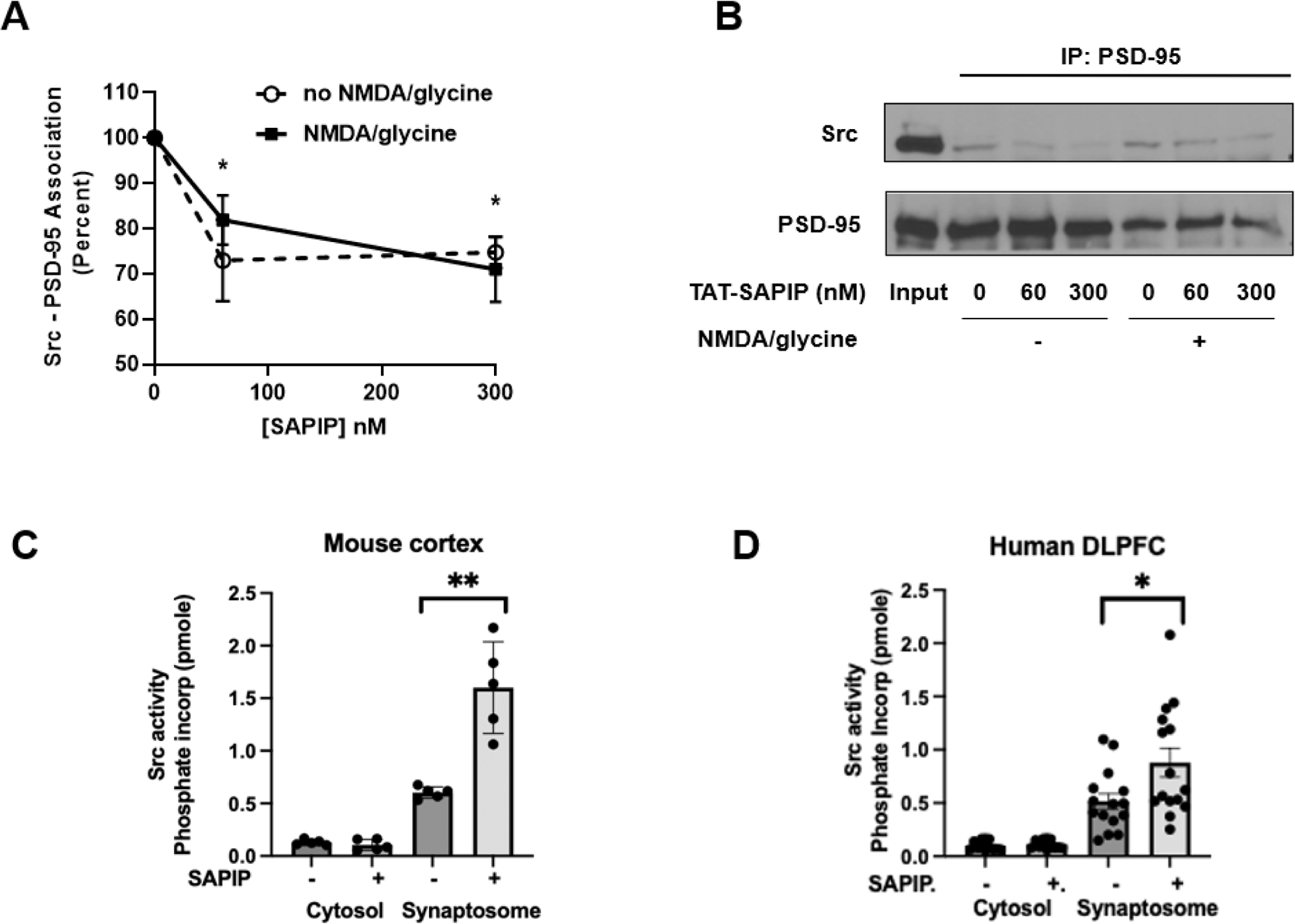
TAT-SAPIP reduces Src-PSD-95 association and enhances Src activity. (A,B) The effects of TAT-SAPIP on Src-PSD-95 association. Rat primary cortical neurons were treated with TAT-SAPIP with or without 10 μM NMDA + 1μM Glycine. Synaptosomal extracts were IPed for PSD-95 and probed for Src. Src-PSD-95 association was significantly decreased in the TAT-SAPIP treatment group (A) Graphic representation, (B) representative blots. (C,D) The effects of TAT-SAPIP on Src activity. Synaptosomes or cytosol from the PFC of mice (C) or from the DLPFC of 15 human subjects (D) were incubated with 200 ng of TAT-SAPIP and Src activity was measured as previously described(*13*). SAPIP significantly increased Src activity in synaptosomes derived from mouse (266.6±32.2% of base line, n=10, p=0.0009) or human PFC (170.0±29.4%, n=30, p=0.024). Bar graphs in C and D show mean and standard error of the mean. Black dots represent individual data points.

The next step was to test if TAT-SAPIP indeed enhances Src activity. We hypothesized that TAT-SAPIP will enhance Src activity in the synapse where Src associates with PSD-95, but not in other intracellular environment devoid of Src-PSD-95 association(*16*). We treated the synaptosomes or cytosol derived from mouse prefrontal cortex with TAT-SAPIP or vehicle and assessed Src activity. Src activity differed significantly between the four groups (F(3,16)=49.95, P<0.001), TAT-SAPIP increased Src activity in the synaptosomes (t(8)=5.088, p=0.0009), while it did not in the cytosol (t(8)= 0.9172, p=0.98) (Figure 1C). Next, we tested this approach in human brain tissues. In postmortem DLPFC of 15 human subjects, (7 healthy and 8 subjects with schizophrenia), Src activity differed significantly between the four groups (F(3,50)=19.20, p<0.001). TAT-SAPIP enhanced Src activity in synaptosomes (t(28)=2.381, p=0.024), but not in cytosol of human DLPFC (t(22)=0.8042, p=0.429) (Figure 1D). Thus, TAT-SAPIP enhances Src activity specifically in the synapse where Src is tonically inhibited by PSD-95, but not in the cytosol.

### TAT-SAPIP increases synaptic NMDAR activity

We then assessed whether TAT-SAPIP can increase synaptic NMDAR activity (Figure 2). We administered TAT-SAPIP to hippocampal slices while recording post-synaptic responses at the Schaeffer collateral–CA1 synapses(*17*). Importantly, we found that TAT-SAPIP had no effect *per se* on the efficacy of synaptic transmission. Excitatory post-synaptic potentials (EPSPs), which are mediated by AMPA receptors, were unaffected by bath applying TAT-SAPIP as assessed in either field or whole-cell patch recordings (Supp Figure 1). To determine the effects of TAT-SAPIP on NMDAR-mediated synaptic responses, we used whole-cell recordings to measure pharmacologically isolated NMDAR excitatory post-synaptic currents (EPSCs). TAT-SAPIP was applied directly through the patch pipette during whole-cell voltage-clamp recordings (Figure 2A). We found that NMDAR EPSCs gradually increased to 161 ± 14% of the basal level (p=0.008, TAT-SAPIP vs no peptide; p=0.002, TAT-SAPIP vs baseline) and remained stable for the duration of the recording (Figure 2A).

**Figure 2.**
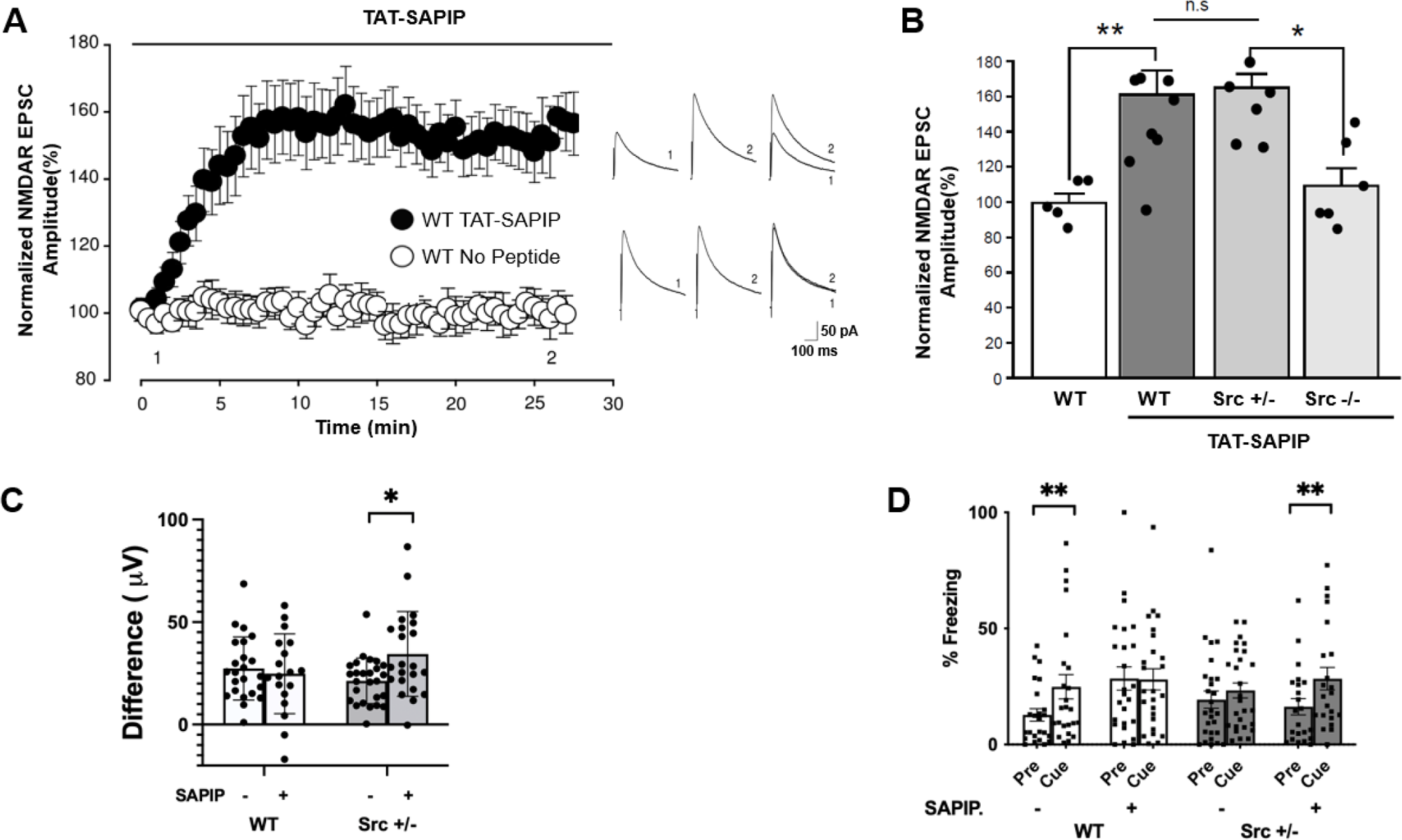
TAT-SAPIP rescues decreased synaptic NMDAR currents and deficits in auditory event related potentials and trace fear conditioning in *Src^+/−^* mice. (A) Scatter plot of NMDAR EPSC peak amplitude over time from WT mice CA1 pyramidal neurons with or without intracellularly applied TAT-SAPIP peptide. Black bar indicates the duration of intracellular peptide application. Right: Representative average NMDAR EPSC traces recorded at membrane potential of +60 mV at the times indicated (1 and 2). B) Histogram representation of normalized NMDAR EPSC amplitude for each group at times denoted by the number 2 in panels A and B. Significant differences were observed in wild-type treated with TAT-SAPIP (161.1±13.8% of baseline, n=10) versus without TAT-SAPIP (100.3±5.1% of baseline, n=5 without peptide) (p=0.009, t-test). Significant differences were also observed between *Src^+/−^* (157.9±7.3% of the baseline, n=8) and *Src^−/−^* (111.6±6.9% of baseline, n=6) groups both treated with TAT-SAPIP (p=0.008, t-test). No differences were observed between WT and *Src^+/−^* when treated with TAT-SAPIP. Black dots represent individual data points. (C) Event-Related Potential response to an oddball stimulus procedure. Difference wave corresponds to the amplitude difference in response to the deviant stimulus minus the standard stimulus. The difference wave of P3a was increased in *Src^+/−^* mice following TAT-SAPIP treatment. Graph depicts mean and SEM while dots represent individual data points. Group sizes were 22 WT vehicle (11 male and 11 female), 26 Src^+/−^ vehicle (14 male and 12 female), 23 WT SAPIP (12 male and 11 female) and 22 Src^+/−^ SAPIP (12 male and 10 female). No effect was observed for sex on any variable. (D) Trace fear conditioning in WT and *Src^+/−^* mice treated with either vehicle or TAT-SAPIP. Graph depicts mean and SEM while dots represent individual data points. WT mice treated with vehicle showed significantly more freezing during the cue relative to pre-cue period (** = p<0.01), demonstrating that they learned to associate the cue with foot shock. In contrast, *Src^+/−^* mice treated with vehicle showed similar levels of freezing during the pre-cue and cue periods, suggesting a failure to learn the association. Treatment with TAT-SAPIP restored trace fear conditioning in *Src^+/−^* mice (** = p<0.01) as indicated with increased freezing during the cue versus pre-cure period. Group sizes were 24 WT vehicle (11 male and 13 female), 27 Src^+/−^ vehicle (12 male and 15 female), 25 WT SAPIP (14 male and 11 female) and 21 Src^+/−^ SAPIP (10 male and 11 female). No effect was observed for sex on any variable.

### TAT-SAPIP is ineffective in *Src^−/−^* mice

To confirm that this increase in NMDAR EPSCs is dependent on Src, TAT-SAPIP was tested in hippocampal slices from homozygous Src knockout (*Src*^−/−^) mice (Figure 2B). In contrast to its effect in wildtype (WT) neurons, TAT-SAPIP failed to affect NMDAR EPSCs in *Src*^−/−^ neurons (109 ± 10% of baseline, p=0.4, TAT-SAPIP vs baseline; p<0.05, *Src*^+/−^ vs WT). Thus, TAT-SAPIP enhancement of NMDAR currents is dependent on the presence of *Src*(*18*). We have previously shown that heterozygous Src (*Src*^+/−^) mice exhibit a subset of behavioral deficits in schizophrenia(*19*). Therefore, we assessed whether TAT-SAPIP can affect synaptic NMDAR activity in these mice. We found that TAT-SAPIP increased NMDAR EPSC amplitude to 165 ± 9.4 % from baseline in *Src^+/−^* neurons (p=0.014, *Src*^+/−^ vs *Src*^−/−^) (Figure 2B). Thus, TAT-SAPIP enhances NMDAR currents in neurons with only a single copy of *Src,* which was used to create a model of decreased Src activity.

### TAT-SAPIP rescues Event-Related Potential and Cognitive deficits in *Src^+/−^* mice

The extent to which behavioral phenotypes associated with the illness are rescued is an important test of a potential therapeutic intervention. Deficits in the P3a auditory event-related potential (ERP) difference waveform are associated with impaired cognitive performance and have been demonstrated in schizophrenia(*20*). To test the potential impact of TAT-SAPIP, we administered this peptide intracerebroventricularly (ICV) for 14 days (Figure 2C). We found that TAT-SAPIP did not alter P3a amplitude difference in WT mice. However, TAT-SAPIP significantly increased P3a amplitude difference in *Src*^+/−^ mice (Figure 2C). These data indicate that TAT-SAPIP enhances the neural circuitry underlying P3a in Src deficient, but not wild-type mice. P3a is a well-established probe for sensory processing abnormalities in schizophrenia(*20*). TAT-SAPIP appears to have restorative effects on the P3a deficits in *Src^+/−^* mice, while disrupting normal function in WT mice (Figure 2C) (gene x treat [F(1,82)=5.51, p=0.021]).

Schizophrenia is characterized by deficits across several cognitive domains subserved by the hippocampus and prefrontal cortex. Trace fear conditioning is a hippocampus-PFC dependent cognitive processes in which the presentation of a cue (CS) and a shock (US) are separated by a stimulus free interval (trace period)(*21, 22*). Successful association of the cue and shock requires retention of the cue across the trace period. A repeated measures ANOVA using percent time freezing during pre-cue versus cue (Cue) periods found a significant three way interaction between gene, treatment and cue [F(1,89)=6.58, p = 0.012). No significant effects were observed for sex. WT mice treated with vehicle showed a significant increase in freezing during the cue compared to pre-cue period (p=0.003) (Figure 2D). In contrast, *Src^+/−^* mice fail to show such difference between the cue and pre-cue (p=0.287), exhibiting a deficit in associating CS and US over a trace period. Chronic ICV TAT-SAPIP restored trace fear conditioning in *Src^+/−^* mice, as indicated by significantly increased freezing to the cue versus pre-cue (p=0.004). However, TAT-SAPIP disrupted trace fear conditioning in WT mice, which showed high freezing during both cue and pre-cue periods (p=0.92) (Figure 2D). Thus, TAT-SAPIP rescued cognitive behavioral deficits associated with Src hypofunction, while disrupting function in wild-type mice. This finding suggests that there is a classical inverted U-shaped response profile with an optimal state that is disrupted by either increased or decreased NMDAR function.

Our findings show that TAT-SAPIP decreases Src-PSD-95 association, increases NMDAR signaling and rescues behavioral and electrophysiological phenotypes associated with NMDAR hypofunction. To test if TAT-SAPIP administration has effects beyond *Src*^+/−^ mice, we then examined mice lacking dysbindin-1, *Sdy*^−/−^ mice, which have multiple neurobiological deficits implicated for schizophrenia, including altered NMDA receptor signaling(*23, 24*). We found that TAT-SAPIP enhances NMDAR EPSCs in *Sdy*+/− mice (Suppl Figure 2). Thus, TAT-SAPIP rescues NMDAR hypofunction in a broader context beyond lack of Src expression.

## Discussion

Data from the current study suggest that blocking the inhibitory effect of PSD-95 on Src may have therapeutic potential for sensory processing abnormalities in schizophrenia. Trace fear conditioning assesses multiple cognitive domains including encoding and retrieval of episodic memory. TAT-SAPIP mediated rescue of deficient trace fear conditioning raises the possibility of an important role for the Src-PSD-95 association in memory and highlights the potential for agents that interfere with protein-protein interactions as a therapeutic approach to reverse highly disabling deficits in neuropsychiatric illnesses.

In the glutamatergic synapses, protein interactions serve as mechanistic substrates by which aberrant expression and function of risk gene products are integrated to precipitate synaptic pathology of schizophrenia. Our results demonstrate that inhibiting the Src-PSD-95 association rescues molecular, electrophysiological, and behavioral phenotypes of NMDAR signaling deficits relevant to schizophrenia. We thus show proof-of-concept that specific protein-protein interactions in the synapse can be leveraged to modify behavioral phenotypes and propose that protein-protein interaction subnetworks of disease relevant pathways be considered as therapeutic targets for neuropsychiatric illnesses.

## Material and Methods

### Study Design

The goal of the current studies was to assess the effectiveness of using TAT-SAPIP to target a protein-protein interaction thought to regulate NMDAR function. It was predicted that TAT-SAPIP would penetrate the cell membrane and increase Src activity. It was predicted that TAT-SAPIP would enhance Src activity in synaptosomes derived from mouse cortex and postmortem DLPFC tissues of human subjects. It was predicted that the effectiveness of TAT-SAPIP would depend directly on Src activity and, as such, TAT-SAPIP would be ineffective in *Src^−/−^* mice. Further, it was predicted that this would lead to increases in NMDAR activity in hippocampal slice tissue. Finally, it was predicted that TAT-SAPIP would rescue ERP and cognitive deficits seen in *Src^+/−^* mice.

Sample sizes are reported in figure captions and were determined based on historical precedence to be the minimal size to detect likely meaningful differences in similar experiments. For P3a and TFC all mice born during the period of the experiment were used. Both experiments were stopped prior to data analysis. No data were excluded from IP, western blot or Src activity assays. No data were excluded from TFC. For EEG experiments mice were removed if the headcap became detached from the skull or if the EEG recording failed to show an expected ERP signal using pre-established criteria developed in the Siegel lab. EEG recordings were assessed for quality by three individuals (O.M., R.E.F, and S.J.S.). For P3a and TFC and Grubbs outlier test was run. One mouse was removed from the P3a experiment due to an excessively high P3a waveform. No mice were removed from the TFC experiment.

Experiments were conducted in both human (postmortem tissue) and mouse subjects. Postmortem experiments were approved by the Institutional Review Board at the University of Pennsylvania. Subjects were prospectively diagnosed by DSM criteria and consents for autopsy were obtained from the next-of-kin or a legal guardian.

Flash-frozen DLPFC tissues from 15 subjects (7 SCZ subjects, and 8 controls) were then obtained from the Penn Brain Bank at the University of Pennsylvania. Details of subjects are included in supplementary table 1. All participants were Caucasian except for one. Postmortem brains were randomly selected from a larger cohort in a relatively balanced combination of control and schizophrenia subjects. Src +/− mice were obtained from Jackson Laboratories on a B6;129S7 background (Jackson labs strain number 002381, B6;S129 Srctm1Sor/J B6;129S7). Mice were between 1 to 6 months of age. All procedures related to ex vivo recordings were performed at the Hospital for Sick Children in accordance the Canadian Council on Animal Care and approved by the Animal Care Committee of the Hospital for Sick Children. All procedures related to in vivo testing were performed at the University of Southern California in accordance with the National Institutes of Health Guide for the Care and Use of Laboratory Animals and approved by the Institutional Animal Care and Use Committee at the University of Southern California.

For in vivo experiments Src+/− and WT mice have a difference in coat color that makes obvious the genotype of each mouse. As such, it is not possible to run a completely blinded experiment. However, experimenters were blinded to the treatment condition of individual mice (SAPIP versus Vehicle). Experimenters conducting IP, western blot and Src activity assays were blinded to the identity of each sample..

### Statistical Analysis

No data were transformed, re-coded, re-scaled or normalized. All in vitro data were analyzed via ANOVA or t-test. In vivo experiments were analyzed via repeated measures ANOVA with time spent freezing during the cue versus pre-cue periods as the repeated dependent variable and sex, genotype and treatment as independent variables. Post hoc tests were conducted using planned comparison t-tests. In all analyses, alpha was set at p<0.05. All statistical tests were two tailed.

### Postmortem brains

Flash-frozen DLPFC tissues from 15 subjects (7 SCZ subjects, and 8 controls), obtained from the Penn Brain Bank at the University of Pennsylvania were used for the study. Approved by the Institutional Review Board at the University of Pennsylvania, subjects were prospectively diagnosed by DSM criteria and consents for autopsy were obtained from the next-of-kin or a legal guardian. Subjects with a history of substance abuse, neurological illnesses, or the need for ventilator support near death were excluded. (Additional details in supplemental data). Detailed demographic data is shown in Suppl Table 1.

### Animals

Src heterozygous C57BL/6 mice were obtained from The Jackson Laboratory (Bar Harbor, Maine), and bred to acquire genotypes *Src^−/−^*, *Src^+/−^* and wild type (WT; *Src^+/+^)* (*25*). Dysbindin-1 (DTNBP1) heterozygous mice were obtained from The Jackson Laboratory. Mice were housed in ventilated cages on a 12:12-h light/dark cycle, with lights on from 7am to 7pm. All *ex vivo* recordings and *in vivo* testing occurred during the light cycle. All procedures related to *ex vivo* recordings were performed at the Hospital for Sick Children in accordance the Canadian Council on Animal Care and approved by the Animal Care Committee of the Hospital for Sick Children. All procedures related to *in vivo* testing were performed at the University of Southern California in accordance with the National Institutes of Health Guide for the Care and Use of Laboratory Animals and approved by the Institutional Animal Care and Use Committee at the University of Southern California.

### Rat Cortical Primary Culture

Primary cultures of rat cortical neurons were obtained from the Mahoney Institute of Neurological Sciences Neuron Culture Service at the Perelman School of Medicine. The neurons were plated at a density of 350,000 cells/mL in Neurobasal media supplemented with B27.

### Preparation of SAPIP

The Src activating PSD-95 inhibitory peptide (or SAPIP) was generated as GST-fusion proteins as described previously(*15*) or as TAT-SAPIP with Tat protein transduction domain (YGRKKRRQRRR) attached to the N-terminal of the SAPIP. GST fusion proteins were expressed in E. coli and purified by GST affinity chromatography (GE) as previously described by having them expressed in bacteria and according to the manufacturer’s instructions. TAT-SAPIP was purified by Nickel affinity chromatography (Qiagen) following the protocols from the manufacturer.

### SAPIP induced activation of Src

A) Rat cortical neurons. Cultured neurons were washed with 1xPBS and treated first with 0, 60 and 300 nM of TAT-SAPIP in Krebs Ringer (KR) solution at 37°C for 15 minutes. This was followed by further incubation in the presence or absence of 10 uM NMDA and 1 uM Glycine at 37°C for 30 more minutes. The cells were collected and processed to obtain synaptosomal fractions for immunoprecipitation and western blotting as described below. B) Mouse or human postmortem brains. Synaptosomes and cytosols were obtained from the PFC of C57BL6 mice or postmortem DLPFC from 15 subjects. Twenty mgms of synaptosomes or cytosol were incubated with 200 nanogms of GST-SAPIP or GST at 4C overnight as previously described.

### Src activity assay

Following the incubation with or without SAPIP, synaptosomes from mouse or human postmortem brains were assayed for Src activity as previously described using Src assay kit (Upstate)(*13*).

### Subcellular Fractionation, Western Blotting and Immunoprecipitation

Synaptosome, synaptic membranes or PSD enriched fractions were isolated from the prefrontal cortex of the mouse or rat cortical cultures by methods from previous studies (*13, 16*). Soluble and insoluble fractions were collected and analyzed by immunoblotting. Synaptic membrane (P2) enriched fractions were immuno-precipitated with anti-PSD-95 (Neuro Mab, mouse monoclonal) and mixed with agarose-conjugated protein A-G beads. Solubilized immunoprecipitates were size separated, and immuno-blotted with anti-PSD-95 (Neuro Mab, mouse monoclonal). Western Blotting was also carried out as indicated in the figures for GluN1/GluN2A (Santa Cruz, goat polyclonal) or Src (Cell Signaling, mouse monoclonal). Signals were detected with ECL (Amersham), developed on X-ray film and quantified using densitometric scanning.

### Hippocampal *Ex Vivo* Slice Electrophysiology

Parasagittal hippocampal slices (300 μm) were prepared in ice-cold ACSF from mice (22-28 days of age) anesthetized (20% (wt/vol) urethane, intraperitoneal (i.p.). Slices were placed in a holding chamber (30 °C) for 40min and then allowed it to passively cool down to room temperature (21 to 22 °C for ≥ 30min) before recording. A single slice was transferred to a recording chamber and superfused with ACSF at 4 ml min^−1^ composed of (in mM) 124 NaCl, 2.5 KCl, 1.25 NaH_2_PO_4_, 2 MgCl_2_, 11 D-glucose, 26 NaHCO_3_ and 2 CaCl_2_ (ACSF chemicals from Sigma-Aldrich) saturated with 95% O_2_ (balance 5% CO_2_) at room temperature, pH 7.40, osmolality 305 mOsm. Synaptic responses were evoked by stimulating Schaffer collateral afferents using bipolar tungsten electrodes located ∼50 μm from the pyramidal cell body layer in CA1. Whole-cell patch-clamp recordings of CA1 pyramidal neurons were carried out using the visualized method (Zeiss Axioskop 2FS microscope). Patch pipettes (4–5 MΩ) solution containing (in mM): Cs gluconate 117, CsCl 10, BAPTA 10, CaClRR_2_RR 1, HEPES 10, ATP-Mg 2, QX-314 10, GTP 0.3 (pH 7.25, osmolality 290 mOsm). Testing stimuli (0.1 ms in duration) were delivered at a frequency of 0.1Hz to evoke synaptic transmission. NMDAR-mediated EPSCs were pharmacological isolated by blockade of AMPA receptors with bath-applied CNQX (10 µM). Bicuculline (10 µM) was included in the bath to block GABARR_A_RR-receptor-mediated transmission. We amplified raw data using a MultiClamp 700B amplifier and a Digidata 1322A acquisition system sampled at 10 KHz and analyzed the data with Clampfit 10.6 (Axon Instruments). Statistical analysis was conducted using SigmaPlot software version 12. The tests used and p values are reported in the respective figure legends.

### Surgery for *In Vivo* Experiments

Mice (8-18 weeks of age) were anesthetized with 1% Isoflurane and implanted with a low-impedance (<5 kΩ, 1000 Hz) stainless steel tripolar electrode (PlasticsOne, Roanoke, VA). Electrodes were cut to a length of 3 mm (positive) and 1 mm (ground and reference) and placed at 1 mm intervals along the sagittal axis. The positive electrode was positioned 1.8mm posterior, 2.65 mm right lateral, and 2.75 mm deep relative to bregma as previously described (*26–28*). A cannula (Alzet Brain Infusion Kit #3, Durect Corporation, Cupertino, CA), was placed 1mm left lateral and 0.5mm posteriorly relative to bregma (Devos and Miller, 2013). The electrode and cannula were secured to the skull with dental cement (Ortho Jet; Lang Dental, Wheeling, IL) and ethyl cyanoacrylate (Elmers, Columbus, OH). The catheter connected to the base of the cannula was driven subcutaneously on the dorsal surface and the implant was closed with a single stitch(*29*). Supplemental warmth was provided post-surgery with the heating pad. A micro-osmotic pump (Alzet Model #1002, Durect Corporation, Cupertino, CA) was primed for a 48-hour period at 37C with TAT-SAPIP peptide dissolved in vehicle to achieve a final dose of 1 ug/kg/day. Osmotic pumps were implanted for 14 days. Animals were monitored throughout the course of the study for weight loss, signs of distress and adverse reactions at the implant site.

### Deviance Detection

Mice were exposed to a series of 19 repetitive standard tones (6 or 9KHz, 85dB) followed by a novel tone (9 or 6KHz, 85dB) using a flip/flop control procedure. Waveforms were corrected relative to baseline at stimulus onset. *Analysis*: Only the standard tone presented immediately before the novel tone was used for analysis to ensure that an equal number of standard and novel tones were used. Subtraction of the standard from the novel produced a difference wave, which was quantified by the peak negative value within the relevant time period of 30–80 msec post-stimulus for MMN and 60 to 200 msec for the P3A.

### Trace Fear Conditioning

Trace fear conditioning took place in rectangular chambers measuring 40cm long, 15 cm wide × 22 cm high (Med Associates, Fairfax, VA). Training consisted of three sets of conditioned stimuli (CS) – unconditioned stimuli (US) pairings. The CS consisted of a 20 second tone (2,700 Hz, 85 dB) while the US was a 2 second 0.6mA foot shock. The CS and US were separated by a period of 20 seconds (trace period). CS-US pairings were presented at 3, 7 and 11 minutes. Twenty-four hours after training, mice were tested for memory of conditioning. To control for possible influence from contextual learning, the conditioning chambers were altered by placing a thin Plexiglas panel on the floor grid and placing an odorant underneath the floor. The CS was again presented at 3, 7 and 11 minutes, during which time freezing was measured (cue). Freezing was also measured 20s prior to CS onset (pre-cue). Freezing behavior was scored in real time digitally (FreezeScan 1.0, Clever Sys Inc) and was quantified as percent time freezing.

## Acknowledgements

Support was provided by R01 MH075916-05 (Hahn/Siegel) and 5P50MH096891-03 (RE Gur). This work was done in part due to a generous gift from Isaac Larian and Family to support psychosis research at USC.

## Notes

### Competing Interest Statement

The authors have declared no competing interest.

## References

1. A. de Bartolomeis, A. Barone, V. Begni, M. A. Riva, Present and future antipsychotic drugs: A systematic review of the putative mechanisms of action for efficacy and a critical appraisal under a translational perspective. Pharmacol Res 176, 106078 (2022).

2. D. T. Balu, The NMDA Receptor and Schizophrenia: From Pathophysiology to Treatment. Adv Pharmacol 76, 351–382 (2016).

3. D. J. Weiner, A. Nadig, K. A. Jagadeesh, K. K. Dey, B. M. Neale, E. B. Robinson, K. J. Karczewski, L. J. O’Connor, Polygenic architecture of rare coding variation across 394,783 exomes. Nature 614, 492–499 (2023).

4. H. Kato, H. Kimura, I. Kushima, N. Takahashi, B. Aleksic, N. Ozaki, The genetic architecture of schizophrenia: review of large-scale genetic studies. J Hum Genet 68, 175–182 (2023).

5. M. J. Gandal, J. R. Haney, N. N. Parikshak, V. Leppa, G. Ramaswami, C. Hartl, A. J. Schork, V. Appadurai, A. Buil, T. M. Werge, C. Liu, K. P. White, S. Horvath, D. H. Geschwind, Shared molecular neuropathology across major psychiatric disorders parallels polygenic overlap. Science 359, 693–697 (2018).

6. V. Trubetskoy, A. F. Pardinas, T. Qi, G. Panagiotaropoulou, S. Awasthi, T. B. Bigdeli, J. Bryois, C. Y. Chen, C. A. Dennison, L. S. Hall, M. Lam, K. Watanabe, O. Frei, T. Ge, J. C. Harwood, F. Koopmans, S. Magnusson, A. L. Richards, J. Sidorenko, Y. Wu, J. Zeng, J. Grove, M. Kim, Z. Li, G. Voloudakis, W. Zhang, M. Adams, I. Agartz, E. G. Atkinson, E. Agerbo, M. Al Eissa, M. Albus, M. Alexander, B. Z. Alizadeh, K. Alptekin, T. D. Als, F. Amin, V. Arolt, M. Arrojo, L. Athanasiu, M. H. Azevedo, S. A. Bacanu, N. J. Bass, M. Begemann, R. A. Belliveau, J. Bene, B. Benyamin, S. E. Bergen, G. Blasi, J. Bobes, S. Bonassi, A. Braun, R. A. Bressan, E. J. Bromet, R. Bruggeman, P. F. Buckley, R. L. Buckner, J. Bybjerg-Grauholm, W. Cahn, M. J. Cairns, M. E. Calkins, V. J. Carr, D. Castle, S. V. Catts, K. D. Chambert, R. C. K. Chan, B. Chaumette, W. Cheng, E. F. C. Cheung, S. A. Chong, D. Cohen, A. Consoli, Q. Cordeiro, J. Costas, C. Curtis, M. Davidson, K. L. Davis, L. de Haan, F. Degenhardt, L. E. DeLisi, D. Demontis, F. Dickerson, D. Dikeos, T. Dinan, S. Djurovic, J. Duan, G. Ducci, F. Dudbridge, J. G. Eriksson, L. Fananas, S. V. Faraone, A. Fiorentino, A. Forstner, J. Frank, N. B. Freimer, M. Fromer, A. Frustaci, A. Gadelha, G. Genovese, E. S. Gershon, M. Giannitelli, I. Giegling, P. Giusti-Rodriguez, S. Godard, J. I. Goldstein, J. Gonzalez Penas, A. Gonzalez-Pinto, S. Gopal, J. Gratten, M. F. Green, T. A. Greenwood, O. Guillin, S. Guloksuz, R. E. Gur, R. C. Gur, B. Gutierrez, E. Hahn, H. Hakonarson, V. Haroutunian, A. M. Hartmann, C. Harvey, C. Hayward, F. A. Henskens, S. Herms, P. Hoffmann, D. P. Howrigan, M. Ikeda, C. Iyegbe, I. Joa, A. Julia, A. K. Kahler, T. Kam-Thong, Y. Kamatani, S. Karachanak-Yankova, O. Kebir, M. C. Keller, B. J. Kelly, A. Khrunin, S. W. Kim, J. Klovins, N. Kondratiev, B. Konte, J. Kraft, M. Kubo, V. Kucinskas, Z. A. Kucinskiene, A. Kusumawardhani, H. Kuzelova-Ptackova, S. Landi, L. C. Lazzeroni, P. H. Lee, S. E. Legge, D. S. Lehrer, R. Lencer, B. Lerer, M. Li, J. Lieberman, G. A. Light, S. Limborska, C. M. Liu, J. Lonnqvist, C. M. Loughland, J. Lubinski, J. J. Luykx, A. Lynham, M. Macek, Jr., A. Mackinnon, P. K. E. Magnusson, B. S. Maher, W. Maier, D. Malaspina, J. Mallet, S. R. Marder, S. Marsal, A. R. Martin, L. Martorell, M. Mattheisen, R. W. McCarley, C. McDonald, J. J. McGrath, H. Medeiros, S. Meier, B. Melegh, I. Melle, R. I. Mesholam-Gately, A. Metspalu, P. T. Michie, L. Milani, V. Milanova, M. Mitjans, E. Molden, E. Molina, M. D. Molto, V. Mondelli, C. Moreno, C. P. Morley, G. Muntane, K. C. Murphy, I. Myin-Germeys, I. Nenadic, G. Nestadt, L. Nikitina-Zake, C. Noto, K. H. Nuechterlein, N. L. O’Brien, F. A. O’Neill, S. Y. Oh, A. Olincy, V. K. Ota, C. Pantelis, G. N. Papadimitriou, M. Parellada, T. Paunio, R. Pellegrino, S. Periyasamy, D. O. Perkins, B. Pfuhlmann, O. Pietilainen, J. Pimm, D. Porteous, J. Powell, D. Quattrone, D. Quested, A. D. Radant, A. Rampino, M. H. Rapaport, A. Rautanen, A. Reichenberg, C. Roe, J. L. Roffman, J. Roth, M. Rothermundt, B. P. F. Rutten, S. Saker-Delye, V. Salomaa, J. Sanjuan, M. L. Santoro, A. Savitz, U. Schall, R. J. Scott, L. J. Seidman, S. I. Sharp, J. Shi, L. J. Siever, E. Sigurdsson, K. Sim, N. Skarabis, P. Slominsky, H. C. So, J. L. Sobell, E. Soderman, H. J. Stain, N. E. Steen, A. A. Steixner-Kumar, E. Stogmann, W. S. Stone, R. E. Straub, F. Streit, E. Strengman, T. S. Stroup, M. Subramaniam, C. A. Sugar, J. Suvisaari, D. M. Svrakic, N. R. Swerdlow, J. P. Szatkiewicz, T. M. T. Ta, A. Takahashi, C. Terao, F. Thibaut, D. Toncheva, P. A. Tooney, S. Torretta, S. Tosato, G. B. Tura, B. I. Turetsky, A. Ucok, A. Vaaler, T. van Amelsvoort, R. van Winkel, J. Veijola, J. Waddington, H. Walter, A. Waterreus, B. T. Webb, M. Weiser, N. M. Williams, S. H. Witt, B. K. Wormley, J. Q. Wu, Z. Xu, R. Yolken, C. C. Zai, W. Zhou, F. Zhu, F. Zimprich, E. C. Atbasoglu, M. Ayub, C. Benner, A. Bertolino, D. W. Black, N. J. Bray, G. Breen, N. G. Buccola, W. F. Byerley, W. J. Chen, C. R. Cloninger, B. Crespo-Facorro, G. Donohoe, R. Freedman, C. Galletly, M. J. Gandal, M. Gennarelli, D. M. Hougaard, H. G. Hwu, A. V. Jablensky, S. A. McCarroll, J. L. Moran, O. Mors, P. B. Mortensen, B. Muller-Myhsok, A. L. Neil, M. Nordentoft, M. T. Pato, T. L. Petryshen, M. Pirinen, A. E. Pulver, T. G. Schulze, J. M. Silverman, J. W. Smoller, E. A. Stahl, D. W. Tsuang, E. Vilella, S. H. Wang, S. Xu, C. Indonesia Schizophrenia, PsychEncode, C. Psychosis Endophenotypes International, G. O. C. Syn, R. Adolfsson, C. Arango, B. T. Baune, S. I. Belangero, A. D. Borglum, D. Braff, E. Bramon, J. D. Buxbaum, D. Campion, J. A. Cervilla, S. Cichon, D. A. Collier, A. Corvin, D. Curtis, M. D. Forti, E. Domenici, H. Ehrenreich, V. Escott-Price, T. Esko, A. H. Fanous, A. Gareeva, M. Gawlik, P. V. Gejman, M. Gill, S. J. Glatt, V. Golimbet, K. S. Hong, C. M. Hultman, S. E. Hyman, N. Iwata, E. G. Jonsson, R. S. Kahn, J. L. Kennedy, E. Khusnutdinova, G. Kirov, J. A. Knowles, M. O. Krebs, C. Laurent-Levinson, J. Lee, T. Lencz, D. F. Levinson, Q. S. Li, J. Liu, A. K. Malhotra, D. Malhotra, A. McIntosh, A. McQuillin, P. R. Menezes, V. A. Morgan, D. W. Morris, B. J. Mowry, R. M. Murray, V. Nimgaonkar, M. M. Nothen, R. A. Ophoff, S. A. Paciga, A. Palotie, C. N. Pato, S. Qin, M. Rietschel, B. P. Riley, M. Rivera, D. Rujescu, M. C. Saka, A. R. Sanders, S. G. Schwab, A. Serretti, P. C. Sham, Y. Shi, D. St Clair, H. Stefansson, K. Stefansson, M. T. Tsuang, J. van Os, M. P. Vawter, D. R. Weinberger, T. Werge, D. B. Wildenauer, X. Yu, W. Yue, P. A. Holmans, A. J. Pocklington, P. Roussos, E. Vassos, M. Verhage, P. M. Visscher, J. Yang, D. Posthuma, O. A. Andreassen, K. S. Kendler, M. J. Owen, N. R. Wray, M. J. Daly, H. Huang, B. M. Neale, P. F. Sullivan, S. Ripke, J. T. R. Walters, M. C. O’Donovan, C. Schizophrenia Working Group of the Psychiatric Genomics, Mapping genomic loci implicates genes and synaptic biology in schizophrenia. Nature 604, 502–508 (2022).

7. T. Singh, T. Poterba, D. Curtis, H. Akil, M. Al Eissa, J. D. Barchas, N. Bass, T. B. Bigdeli, G. Breen, E. J. Bromet, P. F. Buckley, W. E. Bunney, J. Bybjerg-Grauholm, W. F. Byerley, S. B. Chapman, W. J. Chen, C. Churchhouse, N. Craddock, C. M. Cusick, L. DeLisi, S. Dodge, M. A. Escamilla, S. Eskelinen, A. H. Fanous, S. V. Faraone, A. Fiorentino, L. Francioli, S. B. Gabriel, D. Gage, S. A. Gagliano Taliun, A. Ganna, G. Genovese, D. C. Glahn, J. Grove, M. H. Hall, E. Hamalainen, H. O. Heyne, M. Holi, D. M. Hougaard, D. P. Howrigan, H. Huang, H. G. Hwu, R. S. Kahn, H. M. Kang, K. J. Karczewski, G. Kirov, J. A. Knowles, F. S. Lee, D. S. Lehrer, F. Lescai, D. Malaspina, S. R. Marder, S. A. McCarroll, A. M. McIntosh, H. Medeiros, L. Milani, C. P. Morley, D. W. Morris, P. B. Mortensen, R. M. Myers, M. Nordentoft, N. L. O’Brien, A. M. Olivares, D. Ongur, W. H. Ouwehand, D. S. Palmer, T. Paunio, D. Quested, M. H. Rapaport, E. Rees, B. Rollins, F. K. Satterstrom, A. Schatzberg, E. Scolnick, L. J. Scott, S. I. Sharp, P. Sklar, J. W. Smoller, J. L. Sobell, M. Solomonson, E. A. Stahl, C. R. Stevens, J. Suvisaari, G. Tiao, S. J. Watson, N. A. Watts, D. H. Blackwood, A. D. Borglum, B. M. Cohen, A. P. Corvin, T. Esko, N. B. Freimer, S. J. Glatt, C. M. Hultman, A. McQuillin, A. Palotie, C. N. Pato, M. T. Pato, A. E. Pulver, D. St Clair, M. T. Tsuang, M. P. Vawter, J. T. Walters, T. M. Werge, R. A. Ophoff, P. F. Sullivan, M. J. Owen, M. Boehnke, M. C. O’Donovan, B. M. Neale, M. J. Daly, Rare coding variants in ten genes confer substantial risk for schizophrenia. Nature 604, 509–516 (2022).

8. M. Zamanpoor, Schizophrenia in a genomic era: a review from the pathogenesis, genetic and environmental etiology to diagnosis and treatment insights. Psychiatr Genet 30, 1–9 (2020).

9. A. J. Schork, H. Won, V. Appadurai, R. Nudel, M. Gandal, O. Delaneau, M. Revsbech Christiansen, D. M. Hougaard, M. Bækved-Hansen, J. Bybjerg-Grauholm, M. Giørtz Pedersen, E. Agerbo, C. Bøcker Pedersen, B. M. Neale, M. J. Daly, N. R. Wray, M. Nordentoft, O. Mors, A. D. Børglum, P. Bo Mortensen, A. Buil, W. K. Thompson, D. H. Geschwind, T. Werge, A genome-wide association study of shared risk across psychiatric disorders implicates gene regulation during fetal neurodevelopment. Nat Neurosci 22, 353–361 (2019).

10. G. Kirov, A. J. Pocklington, P. Holmans, D. Ivanov, M. Ikeda, D. Ruderfer, J. Moran, K. Chambert, D. Toncheva, L. Georgieva, D. Grozeva, M. Fjodorova, R. Wollerton, E. Rees, I. Nikolov, L. N. van de Lagemaat, A. Bayes, E. Fernandez, P. I. Olason, Y. Bottcher, N. H. Komiyama, M. O. Collins, J. Choudhary, K. Stefansson, H. Stefansson, S. G. Grant, S. Purcell, P. Sklar, M. C. O’Donovan, M. J. Owen, De novo CNV analysis implicates specific abnormalities of postsynaptic signalling complexes in the pathogenesis of schizophrenia. Mol Psychiatry 17, 142–153 (2012).

11. J. Hall, N. J. Bray, Schizophrenia Genomics: Convergence on Synaptic Development, Adult Synaptic Plasticity, or Both? Biol Psychiatry 91, 709–717 (2022).

12. H. Y. Wang, M. L. MacDonald, K. E. Borgmann-Winter, A. Banerjee, P. Sleiman, A. Tom, A. Khan, K. C. Lee, P. Roussos, S. J. Siegel, S. E. Hemby, W. B. Bilker, R. E. Gur, C. G. Hahn, mGluR5 hypofunction is integral to glutamatergic dysregulation in schizophrenia. Mol Psychiatry 25, 750–760 (2020).

13. A. Banerjee, H. Y. Wang, K. E. Borgmann-Winter, M. L. MacDonald, H. Kaprielian, A. Stucky, J. Kvasic, C. Egbujo, R. Ray, K. Talbot, S. E. Hemby, S. J. Siegel, S. E. Arnold, P. Sleiman, X. Chang, H. Hakonarson, R. E. Gur, C. G. Hahn, Src kinase as a mediator of convergent molecular abnormalities leading to NMDAR hypoactivity in schizophrenia. Mol Psychiatry 20, 1091–1100 (2015).

14. M. W. Salter, L. V. Kalia, Src kinases: a hub for NMDA receptor regulation. Nat Rev Neurosci 5, 317–328 (2004).

15. L. V. Kalia, G. M. Pitcher, K. A. Pelkey, M. W. Salter, PSD-95 is a negative regulator of the tyrosine kinase Src in the NMDA receptor complex. EMBO J 25, 4971–4982 (2006).

16. C. G. Hahn, A. Banerjee, M. L. Macdonald, D. S. Cho, J. Kamins, Z. Nie, K. E. Borgmann-Winter, T. Grosser, A. Pizarro, E. Ciccimaro, S. E. Arnold, H. Y. Wang, I. A. Blair, The post-synaptic density of human postmortem brain tissues: an experimental study paradigm for neuropsychiatric illnesses. PLoS One 4, e5251 (2009).

17. H. Li, V. Rajani, L. Han, D. Chung, J. E. Cooke, A. S. Sengar, M. W. Salter, Alternative splicing of GluN1 gates glycine site-dependent nonionotropic signaling by NMDAR receptors. Proc Natl Acad Sci U S A 118, (2021).

18. X. M. Yu, R. Askalan, G. J. Keil, 2nd, M. W. Salter, NMDA channel regulation by channel-associated protein tyrosine kinase Src. Science 275, 674–678 (1997).

19. K. R. Ward, R. E. Featherstone, M. J. Naschek, O. Melnychenko, A. Banerjee, J. Yi, R. L. Gifford, K. E. Borgmann-Winter, M. W. Salter, C. G. Hahn, S. J. Siegel, Src deficient mice demonstrate behavioral and electrophysiological alterations relevant to psychiatric and developmental disease. Prog Neuropsychopharmacol Biol Psychiatry 93, 84–92 (2019).

20. G. A. Light, N. R. Swerdlow, M. L. Thomas, M. E. Calkins, M. F. Green, T. A. Greenwood, R. E. Gur, R. C. Gur, L. C. Lazzeroni, K. H. Nuechterlein, M. Pela, A. D. Radant, L. J. Seidman, R. F. Sharp, L. J. Siever, J. M. Silverman, J. Sprock, W. S. Stone, C. A. Sugar, D. W. Tsuang, M. T. Tsuang, D. L. Braff, B. I. Turetsky, Validation of mismatch negativity and P3a for use in multi-site studies of schizophrenia: characterization of demographic, clinical, cognitive, and functional correlates in COGS-2. Schizophr Res 163, 63–72 (2015).

21. J. D. Raybuck, K. M. Lattal, Bridging the interval: theory and neurobiology of trace conditioning. Behav Processes 101, 103–111 (2014).

22. M. A. Burman, C. A. Simmons, M. Hughes, L. Lei, Developing and validating trace fear conditioning protocols in C57BL/6 mice. J Neurosci Methods 222, 111–117 (2014).

23. T. T. Tang, F. Yang, B. S. Chen, Y. Lu, Y. Ji, K. W. Roche, B. Lu, Dysbindin regulates hippocampal LTP by controlling NMDA receptor surface expression. Proc Natl Acad Sci U S A 106, 21395–21400 (2009).

24. G. C. Carlson, K. Talbot, T. B. Halene, M. J. Gandal, H. A. Kazi, L. Schlosser, Q. H. Phung, R. E. Gur, S. E. Arnold, S. J. Siegel, Dysbindin-1 mutant mice implicate reduced fast-phasic inhibition as a final common disease mechanism in schizophrenia. Proc Natl Acad Sci U S A 108, E962–970 (2011).

25. P. Soriano, C. Montgomery, R. Geske, A. Bradley, Targeted disruption of the c-src proto-oncogene leads to osteopetrosis in mice. Cell 64, 693–702 (1991).

26. R. E. Featherstone, R. Shin, J. H. Kogan, Y. Liang, M. Matsumoto, S. J. Siegel, Mice with subtle reduction of NMDA NR1 receptor subunit expression have a selective decrease in mismatch negativity: Implications for schizophrenia prodromal population. Neurobiol Dis 73, 289–295 (2015).

27. R. E. Featherstone, L. R. Nagy, C. G. Hahn, S. J. Siegel, Juvenile exposure to ketamine causes delayed emergence of EEG abnormalities during adulthood in mice. Drug Alcohol Depend 134, 123–127 (2014).

28. E. N. Billingslea, V. M. Tatard-Leitman, J. Anguiano, C. R. Jutzeler, J. Suh, J. A. Saunders, S. Morita, R. E. Featherstone, P. I. Ortinski, M. J. Gandal, R. Lin, Y. Liang, R. E. Gur, G. C. Carlson, C. G. Hahn, S. J. Siegel, Parvalbumin cell ablation of NMDA-R1 causes increased resting network excitability with associated social and self-care deficits. Neuropsychopharmacology 39, 1603–1613 (2014).

29. L. C. Amann, M. J. Gandal, T. B. Halene, R. S. Ehrlichman, S. L. White, H. S. McCarren, S. J. Siegel, Mouse behavioral endophenotypes for schizophrenia. Brain Res Bull 83, 147–161 (2010).

30. K. Talbot, W. L. Eidem, C. L. Tinsley, M. A. Benson, E. W. Thompson, R. J. Smith, C. G. Hahn, S. J. Siegel, J. Q. Trojanowski, R. E. Gur, D. J. Blake, S. E. Arnold, Dysbindin-1 is reduced in intrinsic, glutamatergic terminals of the hippocampal formation in schizophrenia. J Clin Invest 113, 1353–1363 (2004).

