## Supplemental Data for "Blocking Src-PSD-95 interaction rescues glutamatergic signaling dysregulation in schizophrenia"

### Supplementary Data

**1. SAPIP peptide application does not change baseline synaptic transmission.** To investigate whether enhancement of synaptic NMDAR current by SAPIP peptide would induce potentiation of synaptic strength, we monitored baseline synaptic transmission by whole-cell recording EPSPs at CA3-CA1 synapses while applying SAPIP peptide intracellularly. We found that SAPIP peptide intracellular application did not change baseline synaptic transmission (Suppl Figure 1A). This effect was also confirmed by the observation that CA3-CA1 field EPSPs did not change by bath application of SAPIP peptide up to 60 min (Suppl Figure 1B). Notably, TAT-SAPIP did not affect the current (I)-voltage (V) relationships of NMDARs suggesting that the effects of TAT-SAPI is not via NMDAR gating (Suppl Figure 1C). All measurements were taken from distinct samples.

**2. TAT-SAPIP rescues NMDAR hypoactivity in *Sdy*$^{-/-}$mice.** We examined whether TAT-SAPIP can enhance NMDA receptor function in conditions resulting from dysregulation of genes that were implicated for schizophrenia. Among these is Dysbindin-1, which is decreased in the DLPFC of schizophrenia (*30*). In addition, Dysbindin-1 knock out mice (*Sdy*$^{-/-}$) exhibit NMDAR hypoactivity (*13*) as well as decreased Src activity. Like *Src*$^{-/-}$ mice, *Sdy*$^{-/-}$ mice did not exhibit changes in NMDAR current-voltage characteristics nor did they show any changes in NMDAR-AMPAR ratios, (Suppl Figure 2 A & B). Src mediated GluN activity can be induced by delivering Src kinase activating peptide, EPQ(pY)EEIPIA, intracellularly. EPQ(pY)EEIPIA induced enhancement of NMDAR EPSC amplitude in wild-type, which was much reduced in Sdy$^{-/-}$ mice compared to wild-type (149.6 ± 9.3% vs 196.6 ± 12.2% respectively) (Suppl Figure 2C). This may suggest a decrease in the activatable pool of Src in *Sdy*$^{-/-}$. In the presence of TAT-SAPIP, however, NMDAR EPSC amplitude in *Sdy*$^{-/-}$ mice was to a level similar to that of wild-type (167.9 ± 3.7% vs 161.1 ± 13.8% respectively) (Suppl Figure 1 D). In addition, co-administration of EPQ(pY)EEIPIA and TAT-SAPIP resulted in comparable potentiation of NMDAR EPSC amplitudes in neurons from both *Sdy*$^{-/-}$ and wild-type controls (267.0 ± 39.4% vs 270.8 ± 22.0% respectively) (Suppl Figure 2D). These together suggest that TAT-SAPIP can increase the activatable pool of Src in other conditions where Src activity is attenuated in schizophrenia. All measurements were taken from distinct samples.

**Supplementary Figure legends**

**Suppl Figure 1. SAPIP peptide application does not change baseline synaptic transmission.**

(A) Scatter plot of NMDAR EPSC peak amplitude over time recorded from CA1 pyramidal neurons taken from wild type mice with intracellularly applied TAT-SAPIP peptide. We monitored baseline synaptic transmission by whole-cell recording EPSPs at CA3-CA1 synapses while applying SAPIP peptide intracellularly.

(B) Scatter plot of field EPSPs 0ver time recorded from CA1 pyramidal neurons taken from wild type mice with intracellularly applied TAT-SAPIP peptide.

(C) Scatter plot with representative traces showing the current (I)-voltage (V) relationship and reversal potential of NMDAR EPSCs at the end of each recording in panel A. All measurements were taken from distinct samples.

**Suppl Figure 2. SAPIP rescues NMDAR hypofunction in Sdy$^{-/-}$ mice.**

(A) Scatter plot with representative traces showing the current (I)-voltage (V) relationship and reversal potential of NMDAR EPSCs at Schaeffer collatoral-CA1 synapses from *Sdy*$^{-/-}$ and WT mice. (B) NMDAR versus AMPAR current are not different between WT and *Sdy*$^{-/-}$ mice. Representative traces of AMPAR current (black) and NMDAR current (red) from each genotype are displayed above. Black dots represent individual data points. (C) Scatter plot of NMDAR EPSC peak amplitude recorded with intracellularly applied EPQ(pY)EEIPIA peptide from dysbindin knockout (blue) and wild-type mice (white). Black bar indicates the duration of peptide application in both genotypes. Right: Representative average NMDAR EPSC traces were recorded at membrane potential of +60 mV at the times indicated (1 and 2). (D) Histogram representation of normalized NMDAR EPSC amplitude over time recorded from CA1 pyramidal neurons taken from dysbindin knockout (blue) and wild-type mice (white) with intracellularly applied EPQ(pY)EEIPIA peptide alone (WT 196.6 ± 12.2% of baseline, n = 7; *Sdy*$^{-/-}$ 149.6 ± 9.3 % of baseline, n=11; p = 0.01, Student t-test), TAT-SAPIP peptide alone (WT 161.1 ± 13.8% of baseline, n = 10; *Sdy*$^{-/-}$ 167.9 ± 3.7% of baseline, n=7; p = n.s., Student t-test) or with both EPQ(pY)EEIPIA + TAT-SAPIP peptides (WT 270.8 ± 22.0% of baseline, n = 5; *Sdy*$^{-/-}$ 267.0 ± 39.4% of baseline, n = 5; p = n.s.,

Student t-test). Right: Representative average NMDAR EPSC traces from TAT-SAPIP and EPQ(pY)EEIPIA + TAT-SAPIP treated groups recorded at membrane potential of +60 mV at baseline (#1) versus 25 min (#2). All measurements were taken from distinct samples. Black dots represent individual data points.

Table 1. Demographic Characteristics

| ID | Sex | Age | PMI | Rx | pH | CPZ (mg/Day) | Antipsychotic | Race |
|---|---|---|---|---|---|---|---|---|
| 1 | F | 74 | 3.5 | N | 6.62 | N/A | | c |
| 2 | F | 76 | 9.5 | S | 6.52 | 35 | quetiapine | c |
| 3 | F | 89 | 7 | N | 6.3 | N/A | | AA |
| 4 | F | 76 | 9 | S | 6.71 | 303 | risperidone | c |
| 5 | F | 92 | 5 | N | 6.5 | N/A | | c |
| 6 | F | 88 | 7.5 | S | 6.58 | UnK | UnK | C |
| 7 | F | 90 | 6 | N | 5.98 | N/A | | c |
| 8 | F | 95 | 8.5 | S | 6.77 | 0 | none | c |
| 9 | M | 86 | 7 | N | 6.36 | N/A | | c |
| 10 | M | 82 | 19.5 | S | 6.54 | 815 | haloperidol | c |
| 11 | M | 98 | 15 | N | 6.22 | N/A | | c |
| 12 | M | 89 | 15 | S | 6.42 | 102 | thiothixene | c |
| 13 | M | 69 | 11 | N | 6.49 | N/A | | c |
| 14 | M | 81 | 9 | S | 6.19 | 158 | olanzapine | c |
| 15 | F | 72 | 7 | N | 6.92 | N/A | | c |

Supplemental Figure 1

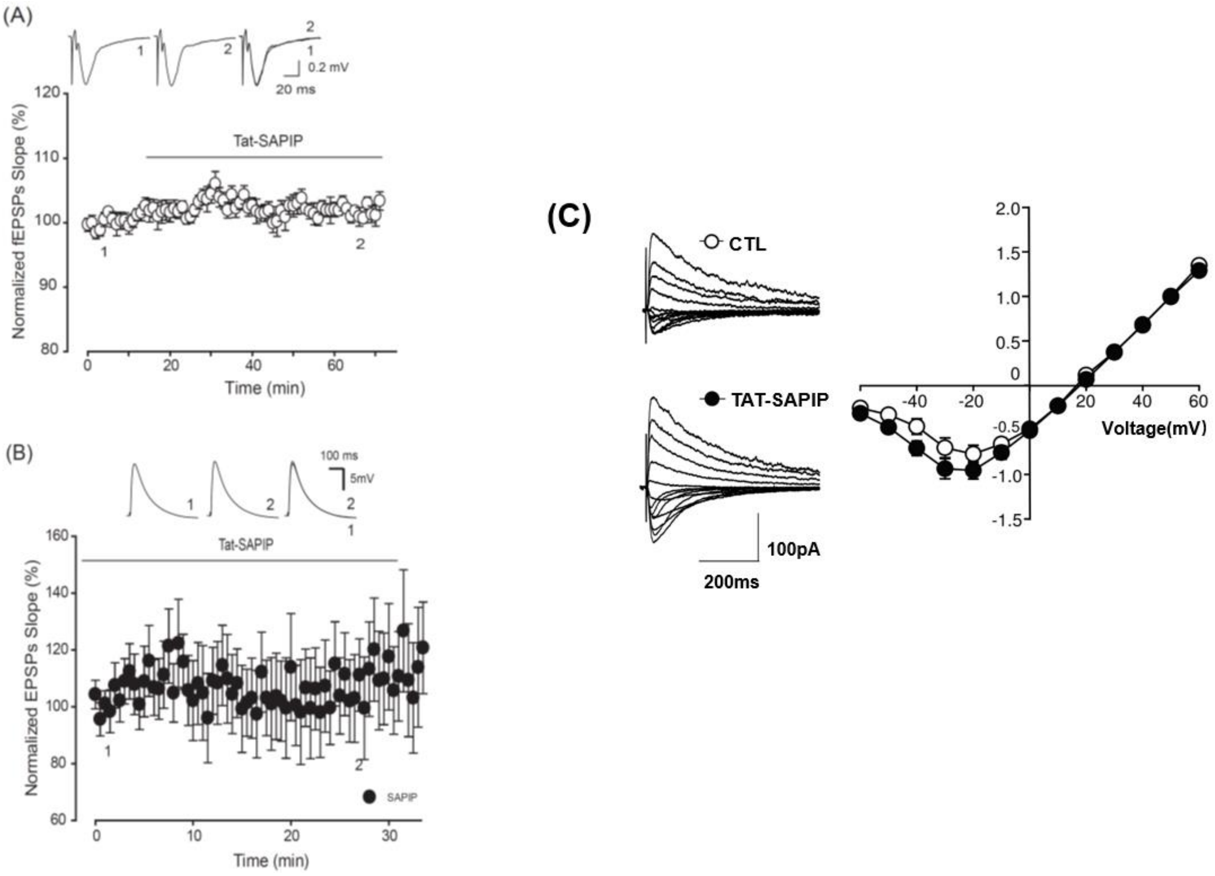

Supplemental Figure 2

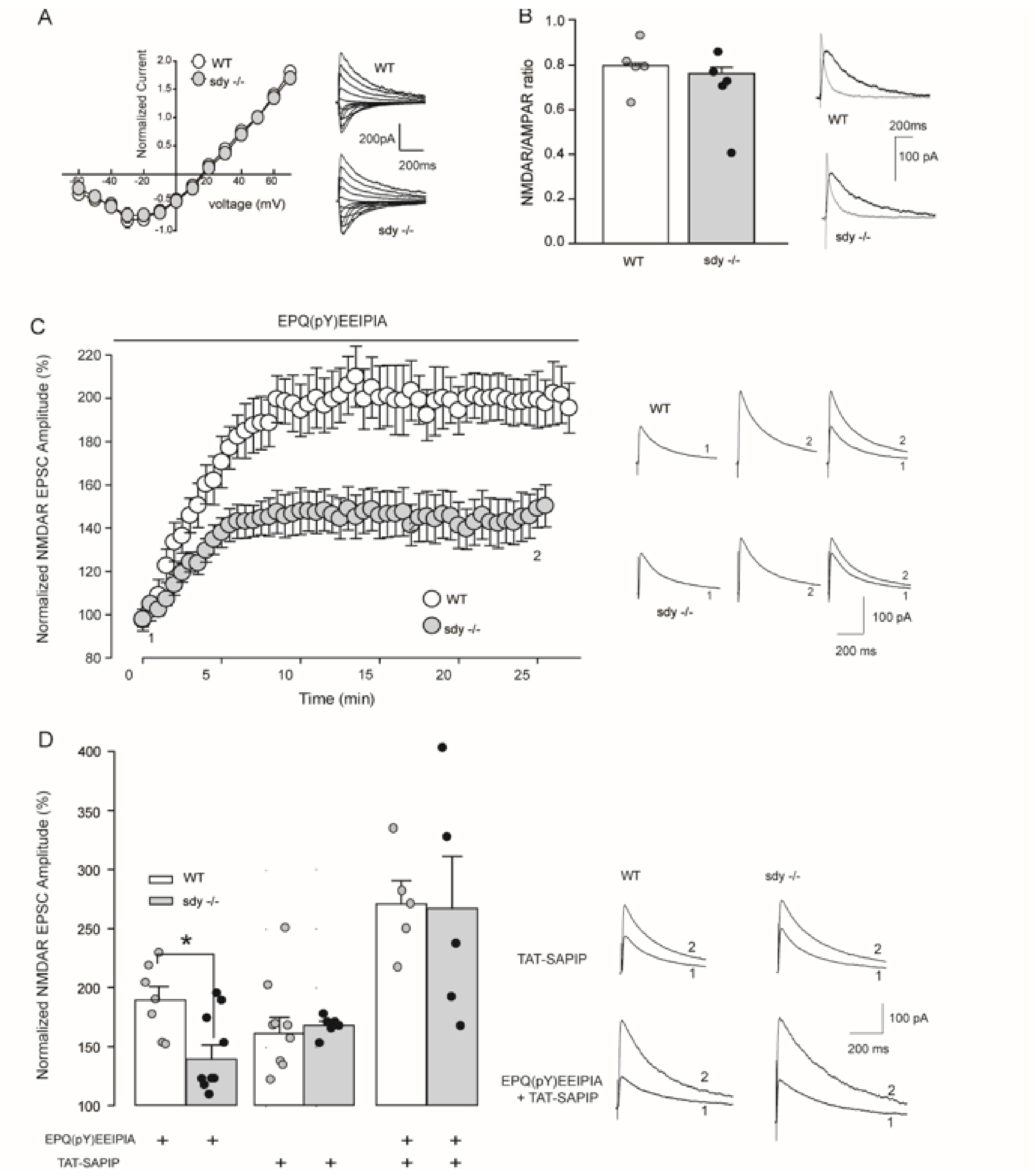
